# Direct visualization of membrane-spanning pores formed by a *Leishmania amazonensis* pore-forming cytolysin, as probed by atomic force microscopy

**DOI:** 10.1101/524686

**Authors:** Thiago Castro-Gomes, J. Mário C. Vilela, Margareth S. Andrade, Paulo S. L. Beirão, Fréderic Frézard, M. Fátima Horta

## Abstract

We have previously shown that *Leishmania amazonensis* produces and secretes a cytolysin that lyses membranes of mammalian cells, including macrophages, its host cell. Using the patch-clamp technique, we have previously demonstrated that the mechanism by which this cytolysin rupture macrophages plasma membrane is by pore formation, which lead us to name it leishporin. While we have characterized leishporin in several aspects, its molecular identity is still unknown. Its behavior suggests that leishporin is, or depend on, a protein, but recent results also suggests that a non-protein molecule is involved in cell lysis. Although the patch-clamp has undeniably revealed that *L. amazonensis* extracts generates pores in macrophages, these structures have not been spotted on cell membranes, which prompted us to several questions: 1) What is the appearance of leishporin-induced pores? Is it similar to that of other described pores? 2) Do these pores physically span lipid bilayers? 4) Are their directly-measured sizes compatible with those previously suggested by patch-clamp? 5) Do these pores fuse with one another, enlarging in size, as suggested by our previous reports? In the present work, we have used two membrane models, erythrocytes and liposomes, to visualize pores induced by the cytolysin on parasite extracts. Leishporin-mediated lysed erythrocytes or liposomes were analyzed by atomic force microscopy (AFM), which allowed us to visualize multiple membrane-spanning pores of variable diameters, ranging from 25 to 230 nm. They do not resemble to protein-formed pores, but rather, to pores made by small molecules such as lipids or peptides, as also visualized by AFM. Our results suggest that the maximum size for individual pores formed by leishporin is around 32 nm, but indicate that they are prone to coalesce, originating large membrane damages that leads to cell collapse, what seems to be a unique property among pore-forming cytolysins.

**Author summary:** One of the mechanisms whereby a cell can be destroyed is by punching holes into their membranes. Through these holes, due to differences in osmolarity between the outside and the inside of a cell, water flows towards the cytoplasm causing plasma membrane ruptures, which damages or lyses cells. We have previously described in the protozoan parasite *Leishmania amazonensis* one of such activities. Using an electrophysiology technique, we have found that parasite extracts lyse cells by making pores on their membranes. However these pores were not directly visualized so far. In this report, using a high-resolution-type scanning microscopy, the atomic force microscopy, we showed in red blood cells membranes and artificial lipid membranes (liposomes) the physical aspect of the pores we described earlier. We observed that these pores are circular-shaped structures with variable diameters, ranging from 25 to 230 nm that span the whole thickness of both types of membranes. We verified that *L. amazonensis* extracts-mediated pores resemble to pores formed by lipids or peptides and not by pores formed by proteins and that they may fuse with one another forming larger holes.

## 1. Introduction

Protozoans of the genus *Leishmania* are intracellular macrophage-dwelling parasites that infect man and other vertebrates. They cause a multifaceted disease known as leishmaniasis, whose different outcomes depend on the more than 20 parasite species, the overall status and the immune response of the host. Vertebrate hosts can be infected by *Leishmania* after being bitten by blood-sucking female sandflies harboring the vertebrate-infective forms, the metacyclic promastigotes. Promastigotes are engulfed by macrophages where they ultimately differentiate into replicative forms, the amastigotes, which settle inside acidic phagolysosomes, aka parasitophorous vacuoles (PV). Eventually, amastigotes leave host cells and can be taken up by neighboring healthy macrophages. The replication of amastigotes, together with their escape from macrophage killing mechanisms and their continuous infectivity to bystander cells, amplifies the infection and, thus, the severity of the disease [1–3]. How amastigotes exit the parasitophorous vacuole and the cell is not yet understood. It has been usually presumed, based on static images only, that unrestricted replication of amastigotes directly causes host cell rupture [4–7] or that amastigotes are released by exocytosis with membrane shriveling, without cell burst [8]. Recently, Real *et al.* (2014) have shown by multidimensional live cell imaging that, *in vitro, L. amazonensis* amastigotes are indeed transferred from infected to healthy macrophages by extrusion of LAMP-rich structures containing the parasite [9]. Whether there is damage to both the parasitophorous vacuole and the plasma membrane is still a controversy.

In previous studies, we have described a cytolytic activity present in *Leishmania amazonensis* amastigostes and promastigostes that functions optimally at 37°C and pH 5.5 (conditions that are found inside the PV), but also at lower temperatures and/or neutral pH [10, 11]. This cytolytic activity has mostly been characterized in *L. amazonensis* [11–13] and *L. guyanensis* [14, 15], but other species such as *L. major* and *L. panamensis* [11] also have it. We have shown that the cytolysin(s) present in *L. amazonensis* extract form transmembrane pores in macrophages, their vertebrate host cell [12], and the molecule(s) responsible for the pore-forming activity has since been designated as leishporin [12, 14, 16]. The binding of leishporin to cell membranes, which occurs directly to membrane phospholipids [13], is a necessary step to form pores and lead to lysis [12].

*L. amazonensis* cytolytic activity seems to result from the removal of a non-covalently linked oligopeptide inhibitor, which can be achieved by proteolytic degradation or simply by dissociation from the active molecule (conditions that are also found inside the PV) [14]. This activation not only improves its lipid-binding efficiency but also endows it with the ability to form pores [15]. Previous results indicated that *L. amazonensis* cytolysin appears to be or depends on a protein present in the membrane fraction of parasite extracts [11, 12] and, indeed, we have identified two genes/proteins that improve *L. major* cytolytic activity. However, we have also identified other non-proteic molecules that may be involved in *L. amazonensis* cytolytic activity (unpublished results). The nature of the molecule(s) we call leishporin is currently under investigation, but is still unknown.

Due to the cytolysin features, we have been postulating that the release of amastigotes from macrophages is not simply a consequence of a physical burden inflicted to the host cells. Rather, it may damage the PV and plasma membranes allowing amastigotes to escape, an event that amplifies the infection, exacerbating the disease and the rate of transmission [11, 12, 16, 17]. The active egress from within the host cell through a variety of strategies, including pore formation, has been described for several intracellular pathogens [16, 18–20].

Pore-formation by promastigote extracts was first suggested by an indirect method, osmotic protection with polyethileneglycol in erythrocytes [10, 11], and first demonstrated by the patch-clamp technique in macrophages [12]. However, we had not yet directly visualized the pores and/or the membrane damage caused by its action. As known from previous works, human erythrocytes [11, 12] and liposomes made from dipalmitoilphosphatidilcholine (DPPC) [13] are lysed after incubation with cytolytically active *L. amazonensis* membrane extracts. Atomic force microscopy (AFM) has been commonly employed to visualize pore-formation produced in biological samples by different pore-forming molecules [19–25]. In the present study, we used the AFM tapping-mode technique in an attempt to visualize cell and artificial membranes damage caused by leishporin. In fact, we were able to visualize the pores we have previously described on the surfaces of lysed erythrocytes and also on lipid films derived from disrupted liposomes that had been treated with *L. amazonensis* membrane extracts. This is the first visual confirmation that the membrane damage caused by *L. amazonensis* pore-forming cytolysin has indeed the appearance of a pore that spans the entire width of the membranes studied. Moreover, a unique property of the cytolysin to form coalescent pores was confirmed.

## 2. Materials and Methods

### 2.1. Parasites

The PH8 (IFLA:PA:67:PH8) strain of *Leishmania (Leishmania) amazonensis,* used throughout this work was provided by Dr. Maria Norma Melo (Departamento de Parasitologia, Universidade Federal de Minas Gerais, Belo Horizonte, Brazil). Parasites were grown in Schnneider medium (Sigma) containing 10% of heat-inactivated fetal bovine serum (GIBCO) and 50 mg/mL of gentamicin (Sigma). Four-day cultured parasites were washed three times with PBS, centrifuged at 1000g and kept at -80°C until required.

### 2.2. Membrane-rich soluble extracts preparation

Parasites were resuspended in borate buffer 50 mM pH 7.0 to a density of 2 x 10^9^ in 1 mL and subjected to five cycles of freeze-and-thaw. After nuclei and debris sedimentation at 1000g for 10 min, the supernatant (crude extracts) was centrifuged for 1 hour at 16.000g. The membrane-rich pellet was resuspended to 1 mL in borate buffer containing 0.4% CHAPS (3[(3-cholamidopropyl)dimethyl-ammonium]-2-hydroxy-propanesulfonate) (a sublytic concentration for erythrocytes) [14] and kept for 1 h on ice with occasional agitation. The suspension was centrifuged at 100,000g for 1 h at 4°C, to remove insoluble material, and the supernatant, corresponding to the solubilized membrane molecules, was collected and referred to as membrane-rich soluble extracts (mExt). Typical preparations contained 1.5 mg/mL protein.

### 2.3. Hemolytic assay

The cytolytic activity of parasite mExt was assessed by a hemolytic assay using human erythrocytes (HuE) as targets, as previously described [11]. Briefly, in 96 round-bottomed well microplates, 5 x 10^6^ HuE in 200 μL of assay buffer (20 mM acetate buffer, 150 mM NaCl, pH 5.5) were incubated with 10 μl of serially diluted fresh or boiled (heat-inactivated - hi) mExt heated. After 30 min at 37°C, the microplates were centrifuged for 10 min at 500g· and hemolysis was quantified by measuring the hemoglobin released, as determined by reading the absorbance of the supernatant at 414 nm. The percentage of lysis was determined in relation to total lysis, obtained by incubation of the same number of erythrocytes with 10 μL of 0.25% Triton X-100. The hemolytic activity was reported as dilution factor (x axis) and *%* lysis (y axis).

### 2.4. Liposome preparation

Liposomes were made of 30 mM DPPC (Lα–dipalmitoylphosphatidylcholine) or DOPC (dioleylphosphatidylcholine). The lipid were dissolved in chloroform in a round bottom flask containing glass beads and connected to a rotatory evaporator at 50°C to evaporate the solvent and form a lipid film. The lipid film was hydrated with PBS buffer to the final lipid concentration of 30 mM. The suspension obtained was subjected to ten cycles of freeze and thaw and extruded trough a polycarbonate membrane of 0.2 μm. The liposomes obtained by this method contain a single lipid bilayer (unilamellar vesicles).

### 2.5. Liposome lysis assay

Ten microliters of calcein-loaded DPPC or DOPC liposomes were diluted in 90 μl of acetate buffer, pH 5.5, and incubated at 0°C or at 37°C with fresh or hi mExt. Twenty-μl aliquots were collected at 5- or 10-min interval and diluted in 2 ml acetate buffer in a quartz cuvette before reading the fluorescence (wavelengths: exciting - 490 nm; emission: 515 nm) (Varian Cary Eclipse Fluorimeter). Lysis was reported as the percentage of total lysis, obtained after addition of 5 μl of 4% CHAPS.

### 2.6. Erythrocytes and lipid film preparation for AFM analysis

In order to visualize the damage caused by leishporin on erythrocytes or liposomes, we incubated cells (2 x 10^6^) or liposomes (20 μl of the suspension described in sections 2.3 and 2.4) with fresh or hi mExt (1:10 or 1:20 depending on the activity of the extract in 1.6 mL (cells) or 200 μl (vesicles) in acetate buffer with a sub-lytic concentration of CHAPS used for extracts preparation on ice (at 0°C to 4°C, leishporin binds to membranes but not form pores, which occurs at higher temperatures from about 20°C to 37°C) for 30 minutes [12, 13]. Cells or liposomes were then centrifuged (10 min at 500g and 10 min 15,000 x g, respectively) and washed three was repeated three to five times to oversupply the membranes with the cytolysin (in general, until no hemolytic activity was detected in the supernatant).

Lipid film preparation from both DPPC and DOPC liposomes was adapted from Puntheeranurak *et al.* (2005) [26] and Richter *et al.* (2006) [27]. After final sedimentation, cells and vesicles were reconstituted to 100 μL and 20 μL, respectively, with acetate buffer then incubated at 37° C for 30 minutes for proper action of the cytolysin. The suspensions containing all lysed cells or liposomes were placed on fresh cleaved mica and dried at 37°C. Once dried, the piece was gently washed to remove salt excess and unfixed liposomes, and dried again. With this treatment liposomes tend to become flat planar and form a homogeneous lipid film [26, 27].

### 2.7. AFM analysis of lysed erythrocytes and lipid films made of lysed liposomes

The cell membranes and the lipid film obtained from DPPC or DOPC liposomes were analyzed by AFM - Tapping Mode (Digital Instruments). AFM images were registered at room temperature (19-23°C) in air with a Dimension 3100 equipment and NanoScope IIIa controller, from Digital Instruments. Tapping mode imaging was performed using silicon cantilevers (AFM/tapping point probes) (Nanosensors), with resonance frequency of 75-98 kHz > and spring constant of 3.0 - 7.1 N/m with images performed with a scan rate of 1 Hz. The set point amplitude was held as high as possible to avoid the influence of the probe pressure on the topographic contrast and to avoid damage of sample surface. Topographic profiles were analyzed by Nanoscope III software (Digital Instruments) and, when necessary, we removed image interference using the Flatten Function in order zero, which do not affect analysis reliability. We analyzed at least ten flat homogeneous fields for each treatment.

All experiments in this work were repeated at least 3 times and all data correspond to a typical result.

## 3. Results

### 3.1. Leishporin-mediated erythrocyte and liposome lysis

We have already shown that, once incubated with promastigotes mExt, erythrocytes and DPPC liposomes are lysed by leishporin, and that boiling mExt inactivates this pore-forming activity [11, 13]. We have also shown that the cytolysin binds directly to the target membranes phospholipids in a cholesterol-independent manner [13]. Here we also show that, interestingly, unlike DPPC liposomes, DOPC liposomes are totally resistant to leishporin-mediated lysis, despite their ability to bind the cytolysin [13]. Figure 1 shows typical lysis kinetic curves of HuE (Fig. 1A) and DPPC liposomes (Fig. 1B) treated with active or heat-inactivated (hi) mExt and DOPC liposomes (Fig. 1C) treated with active mExt during the indicated periods of time. We then used these targets and conditions to visualize the membrane damage caused by leishporin or as negative controls using AFM.

**Figure 1.**
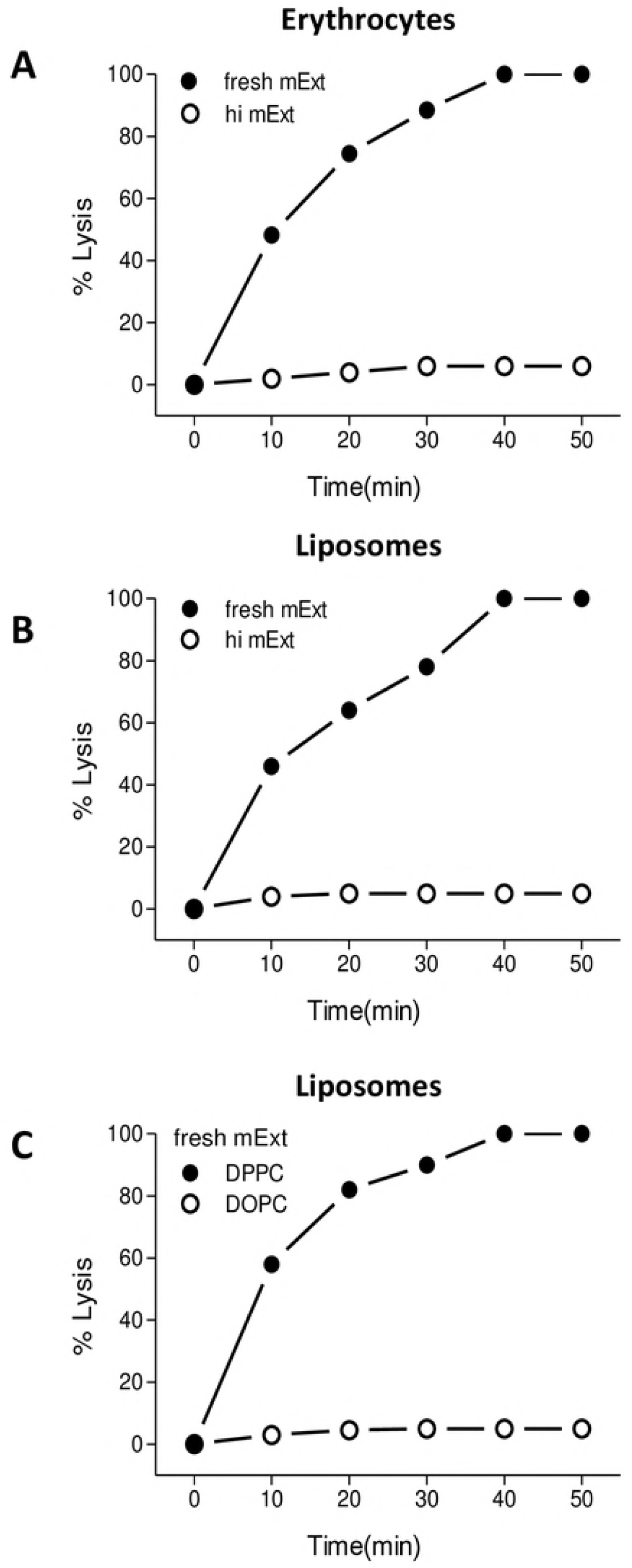
Cytolytic activity of leishporin to erythrocytes and liposomes. Human erythrocytes or calcein-loaded liposomes were incubated with fresh or heat inactivated *L. amazonensis* membrane extract (mExt) at 37° C. To quantify lysis, hemoglobin or calcein released was measured as described in section 2. A-Lysis of human erythrocytes by fresh (dark circles) or heat inactivated (empty circles) mExt. B-Lysis of DPPC liposomes by fresh (dark circles) or heat inactivated (empty circles) mExt. C-Lysis of DPPC (dark circles) and DOPC (empty circles) liposomes by fresh mExt.

### 3.2. Leishporin-mediated pore-formation on erythrocytes

The aspect of HuE treated with fresh or hi mExt, as pictured by AFM, is shown in figure 2. Intact HuE incubated with hi mExt are visualized with its typical size (~6-8 μm) and concavity in figure 2A, whereas leishporin-lysed HuE are shown in figure 2B as cell ghosts. Typical images of their membranes are shown at a higher magnification in figures 2C, 2D, 2E and 2F). Intact HuE (Fig. 2C and 2E) show their irregular, but homogeneous surfaces. A thorough scanning through several fields revealed that the deepest depressions observed are about 3.5 nm deep, indicating that these depressions do not cross the membrane. Examples of those measures are shown in figure 2G in the same field as in figure 2C and 2E. In a striking contrast, the scanned images of the ghost membranes (Fig. 2 D and F), show a totally heterogeneous surface: part resembling to intact cells and part with the presence of many individual circular pore-shaped structures, most of them traversing the whole membrane thickness (blue arrows) (see measurements below). Lesions similar to pores in coalescence (two or more individual pores), frequently presenting borders reminiscent of the original circular structure are also quite distinguishable (yellow arrows). When zoomed in (Fig 2H), the circular edges of these larger structures (yellow arrows) are very clear, visibly indicating the union of two or more circular pores. The topographic analysis of the coalescent structure emphasized in the square in figure 2D is shown in figure 2H, in which two vertical distances were measured (7.7 nm and 5.9 nm), deep enough to cross the plasma membrane, whose thickness is stated to be from 4 to 10 nm [28, 29].

**Figure 2.**
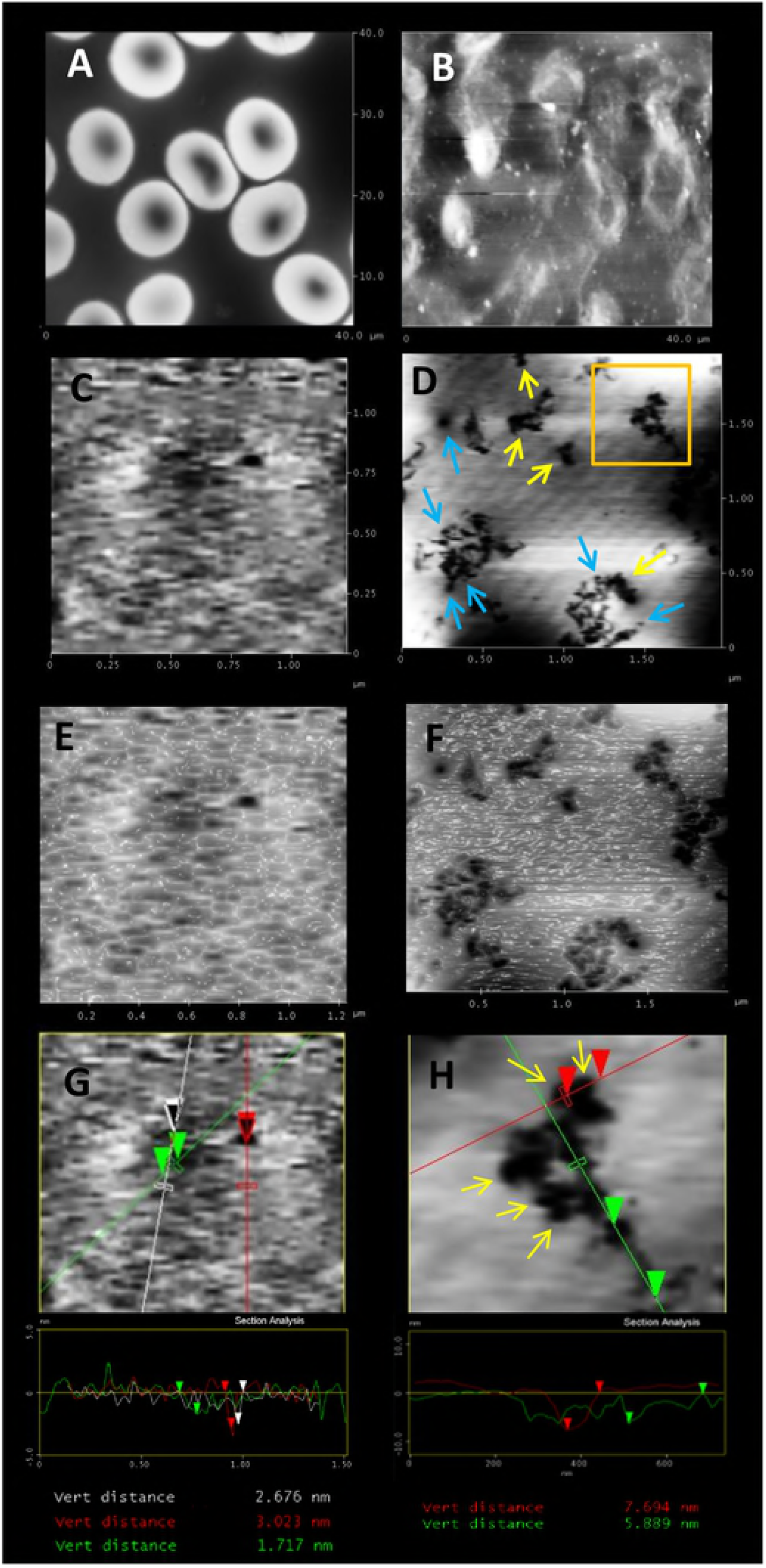
AFM analysis of erythrocytes lysed by leishporin. Erythrocytes were incubated with active mExt, immobilized on mica slices and analyzed by tapping mode AFM. A-Untreated erythrocytes. B-Erythrocytes lysed by mExt (cell ghosts). C-Topological aspect of the plasma membrane of an untreated erythrocyte. D-Topological aspect of the plasma membrane of an erythrocyte lysed by mExt - the image shows several individual (blue arrows) and coalescent (yellow arrows) pores. E, F - 3D views of images shown in C and D, respectively. G, H - Topographic analysis of the plasma membrane regions crossed by the red and green lines in the image shown in C or the zoomed in region defined by the square in the image shown in D, respectively. The measurements correspond to the vertical distance (depth) of each region analyzed.

Another similar experiment, shown in figure 3 (A-C), emphasizes three pores whose depths were around 8.6 nm, 8.2 nm and 7.3 nm (Fig.3 C). In figure 3 we can also see the 3D lateral view of the same field, in which diameter of the larger pore has approximately 50 nm (Fig 3C).

**Figure 3.**
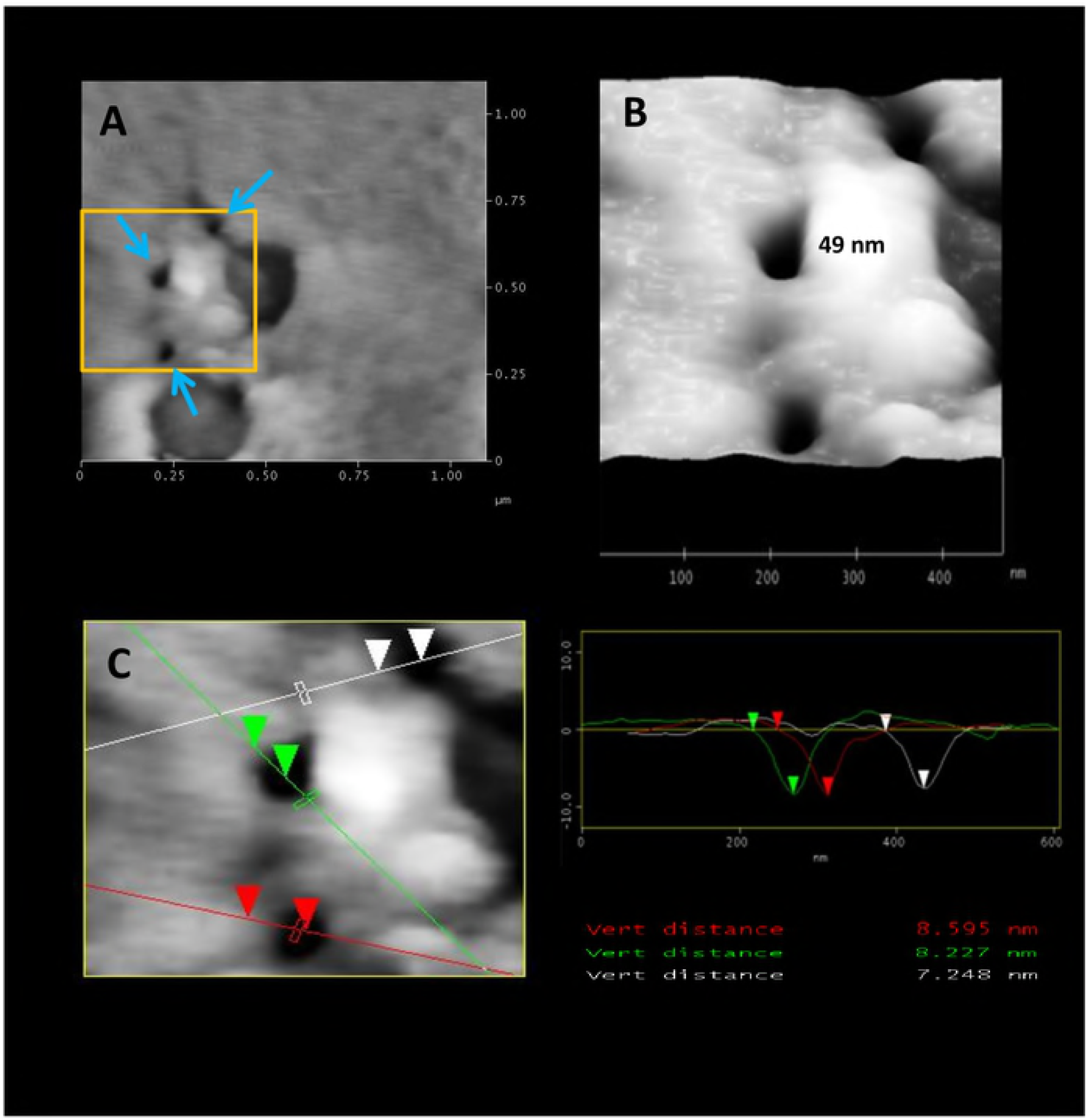
AFM analysis of membrane-spanning pores on erythrocyte lysed by leishporin. The membrane of an erythrocyte lysed by *L. amazonensis* mExt was analyzed in detail by tapping mode AFM. A-Visualization of three individual pores (blue arrows) on the membrane of an erythrocyte lysed by mExt. B-3D view of the zoomed in region defined by the square in the image shown in A. C-Topographic analysis of the regions crossed by the red, white and green lines in the image shown in B. The measurements correspond to the vertical distance (depth) of each pore.

### 3.3. Leishporin-mediated pore-formation on liposomes

Previous results have shown that, in order to lyse cells, leishporin must directly bind to membrane phospholipids. This is clear from experiment shown in figure 1 and from a previous report [13] showing that leishporin is also able to lyse DPPC liposomes, although the cytolysin does not lyse DOPC liposomes (Fig. 1). To investigate whether similar pore-shaped structures were also found in liposomes lysed with mExt, DPPC and DOPC (used as negative controls), liposomes were incubated with cytolytic fresh or hi mExt, as described in section 2, in concentrations sufficient to oversupply the vesicles with the pore-forming molecules. Susceptibility of liposomes to lysis, as seen in figure 1, correlates with the aspect visualized by AFM, represented in figure 4. DOPC incubated with fresh mExt and DPPC incubated with hi mExt presented no signs of disruption (Figs. 4 A and B, respectively). In contrast, DPPC liposomes, incubated with fresh mExt presented many individual (blue arrows) and coalescent pores (yellow arrows) (Fig. 4C), similar to the images from leishporin-lysed erythrocytes. We can see that the depressions/irregularities observed in DOPC or hi-treated DPPC (Figs. 4A and B, respectively), as compared through the scale bars, are quite different from the pores seen in mExt-treated DPPC (Fig. 4C). Figure 4D shows the topographic analysis of the same image shown in figure 4C, in which the some pores measured 7,1 nm, 7,5 nm and 8.2 nm. In DOPC or hi mExt-treated DPPC lipid films all depressions measured have 3.5 nm or less. The absence of pores in DOPC lipid films or DPPC lipid films that have been treated with hi mExt emphasizes that pores only occurs when lysis occurs.

**Figure 4.**
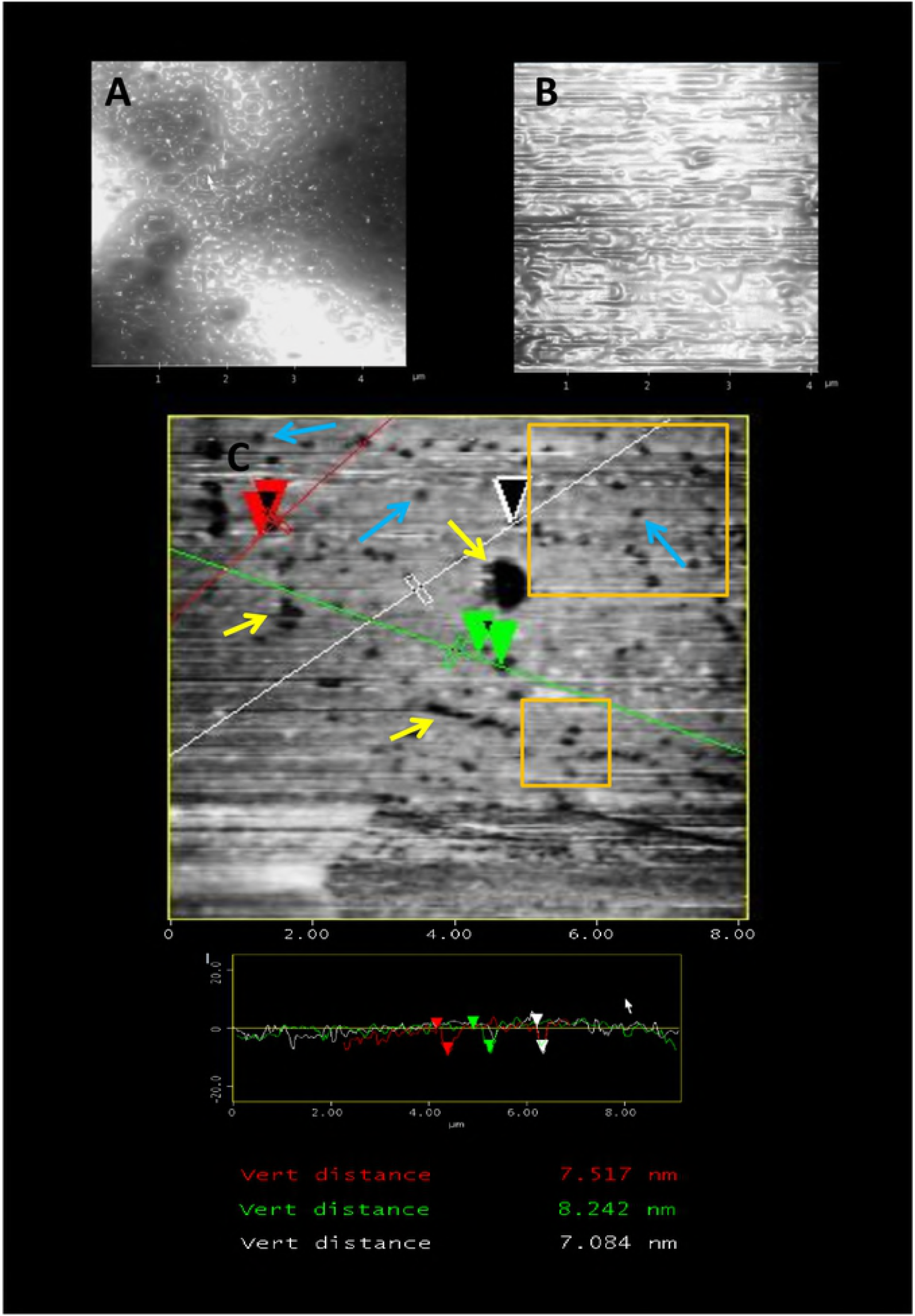
AFM analysis of lipid films made of liposomes lysed by leishporin. Liposomes of each indicated composition were incubated with mExt at 4° C – a condition that allows for cytolysin binding but not for pore formation – washed and then incubated at 37° C to allow for pore formation. Lipid films were prepared with the harvested vesicles and analyzed by AFM. A-Lipid film made of DOPC liposomes (leishporin-insensitive) treated with fresh mExt. B-Lipid film made of DPPC liposomes (leishporin-sensitive) treated with heat-inactivated mExt. C-Lipid film made of DPPC liposomes previously lysed by fresh mExt – the image shows several individual (blue arrows) or coalescent (yellow arrows) pores and the topographic analysis of three individual pores crossed by the red, white and green lines. The measurements correspond to the vertical distance (depth) of each pore analyzed. The areas delimitated by the squares correspond to the zoomed in region shown in Figure 5.

Figure 5A and B (panel 1) represent zoomed in pictures of the larger and smaller square of figure 4D, respectively, where we can see an almost coalescent 2-pores, with the diameter of around 150 nm (Fig. 5B). The topographic profile (Fig. 5C and D (panel 1)) showed that the depths of some other pores, around 6.4 nm, 7.7 nm, 7.8 nm and 8.3 nm, are about the expected thickness of DPPC lipid films, indicating that the pores had traversed the whole extension of the bilayer. Figure 5E (panel 1) shows the 3D lateral view of the two pores of the figures 5B and D (panel 1). One of the pores measured 150 nm in diameter (Fig. 5B, D and E).

**Figure 5.**
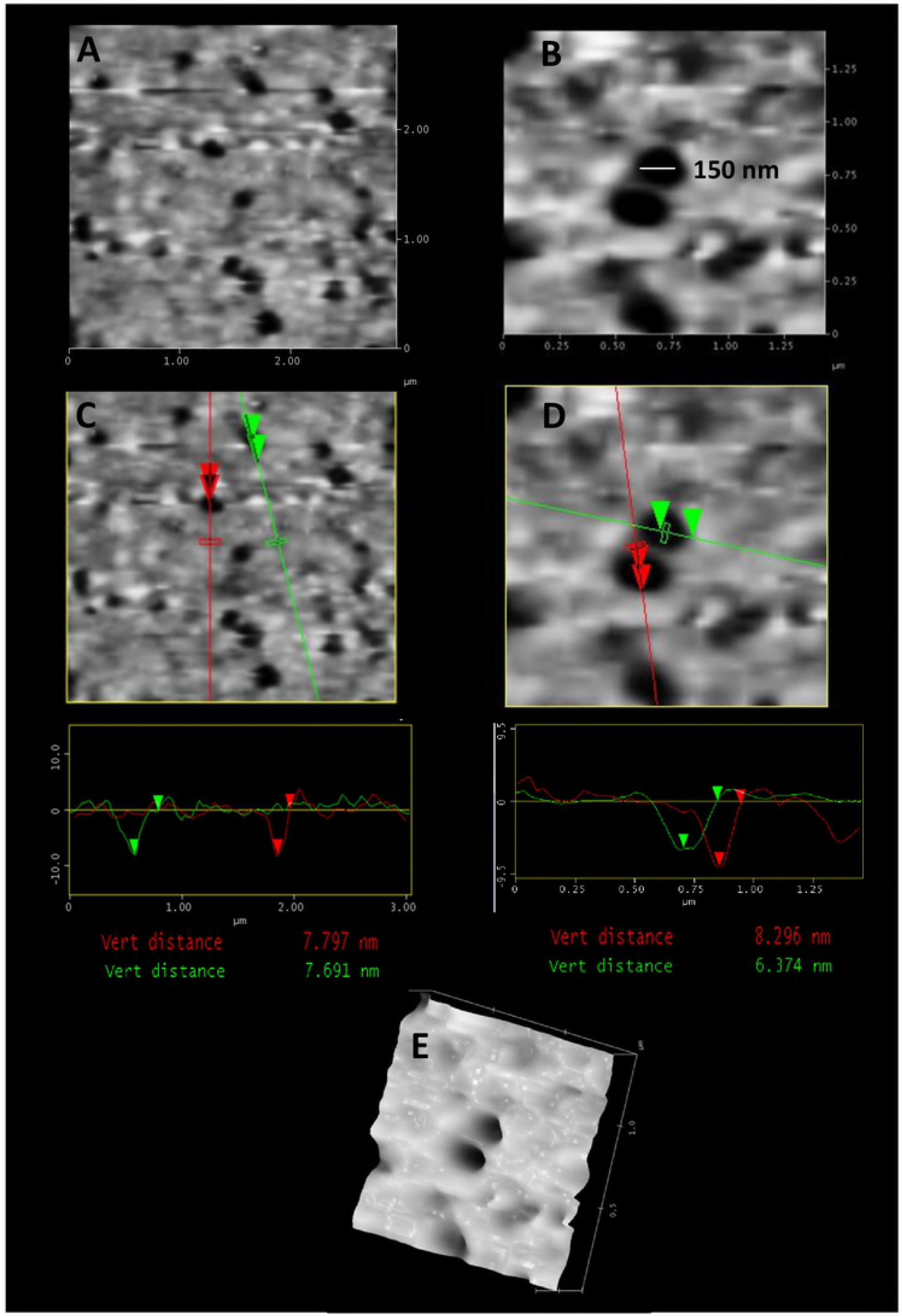
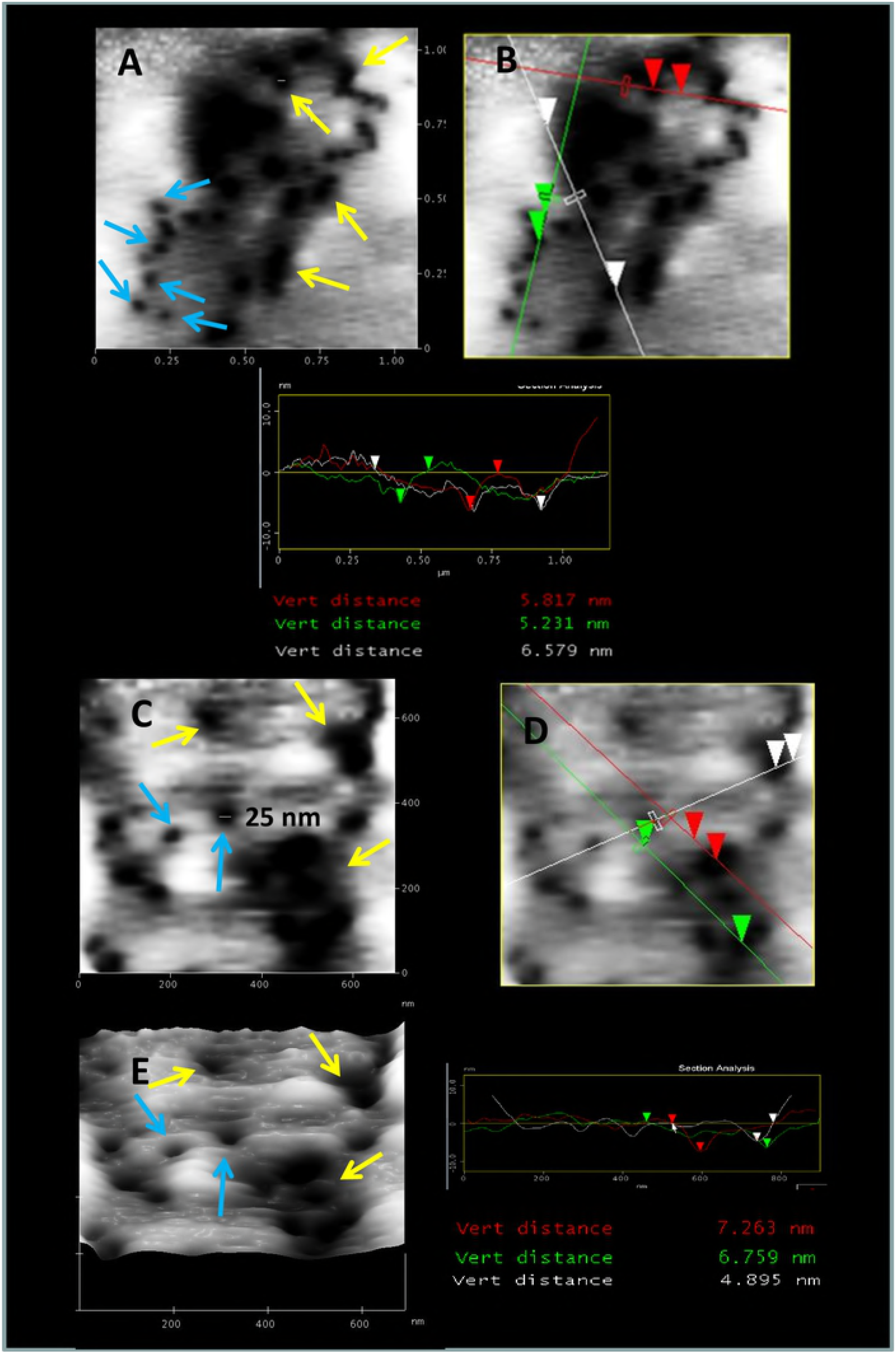
AFM analysis of membrane-spanning pores on lipid films made of DPPC-liposomes lysed by leishporin. DPPC lipid films made of liposomes lysed by mExt were analyzed by AFM. **Panel 1** - Two different areas in images shown in Fig. 4 are zoomed in. A and B - Top view. C-D Topographic analysis of individual pores crossed by the red and green lines. The measures correspond to the vertical distance (depth) of each pore analyzed. E-3D lateral view of the image shown in D. **Panel 2** - Images of a different experiment. A and C - Top view showing individual (blue arrows) and coalescent (yellow arrows) pores. B and D - Topographic analysis of the regions crossed by the red and green lines. The measures correspond to the vertical distance (depth) of each pore analyzed. E-Lateral view in 3D of the area shown in C.

Other fields from different experiments with mExt-lysed DPPC are zoomed in, in figure 5 (panel 2), where individual pores (blue arrows) or coalescent pores (yellow arrows) are also shown (Fig. 5A and C (panel 2)). Figures 5B and D (panel 2) show the depths measurements of some of the pores. In these particular measurements the depths were around 5.2 nm, 5.8 nm, 6.6 nm (Fig. 5B (panel 2)), 4.9 nm, 6.8 nm and 7.3 nm (Fig. 5D (panel 2)). Figure 5E (panel 2) shows the 3D lateral view of figures 5C or D (panel 2).

### 3.4. Frequency of pores dimensions: diameter and thickness

Many of the observed pores were measured to obtain their diameter and depth. Diameter was assumed as the horizontal distance between two points at the half of the vertical distance. Depth was assumed as the vertical distance between the surface line and the deepest point of the valley. Figure 6 shows the frequency of these two parameters. As we can notice, the topographic analysis of all pores measured in erythrocytes showed a vertical distance (thickness) ranging from 4.1 and 8.0 nm measured from 45 clearly defined pores and a few coalescent ones, indicating that all membrane damages crossed the plasma membrane. Around 80% of the pores measured around 5 to 7 nm in depth. The diameters of all pores measured ranged between 32 and 125 nm, including coalescent ones, but most (~70%) measured 45-65 nm (Fig. 6A).

**Figure 6.**
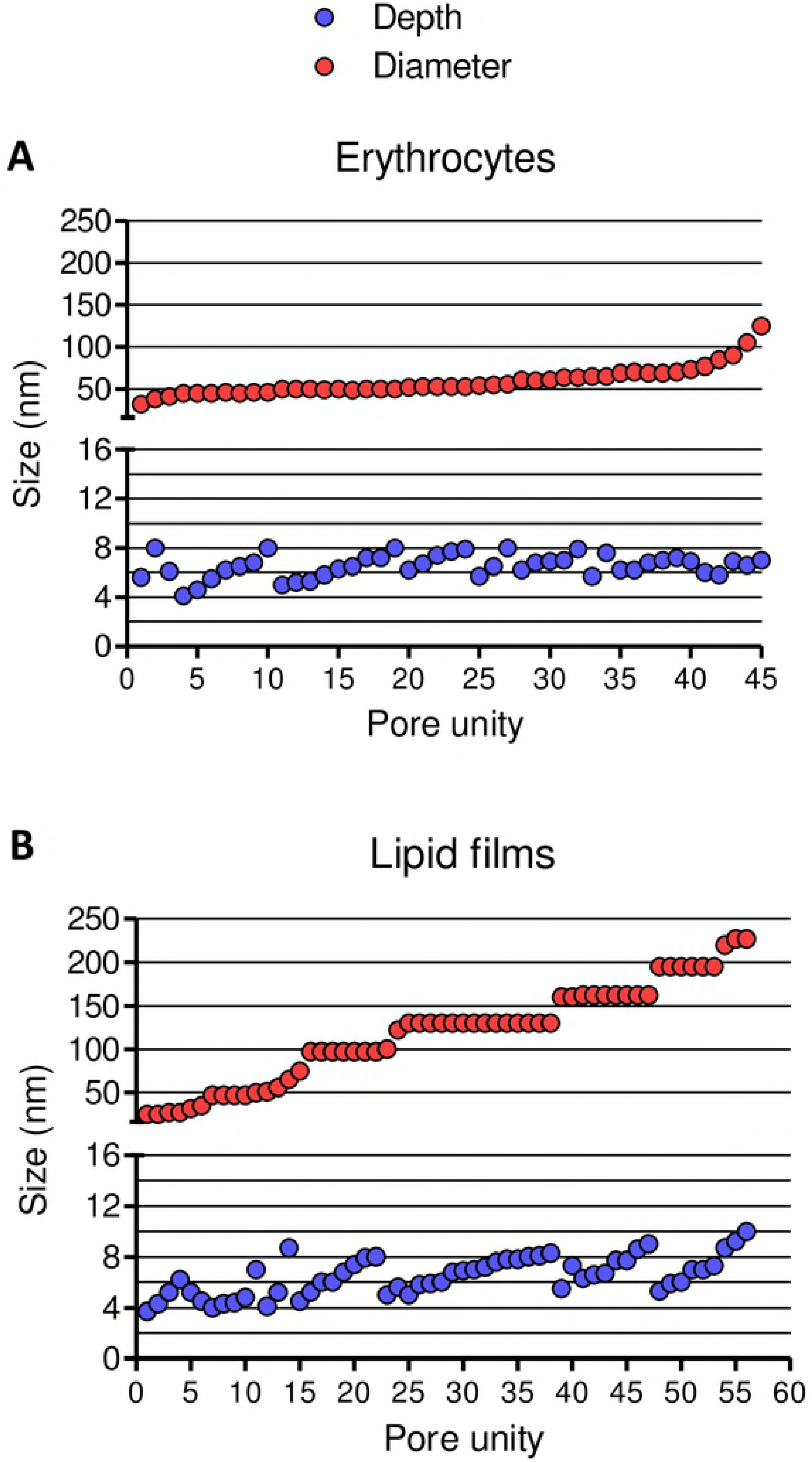
Frequency of dimensions (diameter and thickness) of leishporin membrane-spanning pores on erythrocytes and DPPC lipid films. Dispersion pattern of the dimensions of pores on the membrane of erythrocytes (A) or on lipid films made of DPPC liposomes (B) lysed by mExt. Vertical distances were taken between the surface line and the deepest point observed at the center of the pore. The diameter was defined as the measure of the horizontal distance between two points taken at the half of the vertical distance observed for each pore. Red circles = pore diameter, blue circles = pore depth.

In liposomes, depths varied from 4 nm to 10 nm and, like in erythrocytes, this indicates that pores have traversed the membrane. However, around 80% of pores measured around 5 to 8 nm in depth. Individual or coalescent pores measured from 25 nm to 227 nm, but most (~60%) ranged from 97-162 nm (Fig. 6B). Remarkably, and different from what occurred in erythrocytes, diameters of most pores were multiples of about 32.5 nm, particularly those with diameter having 97 nm or more. Actual measurements of majority of those pores were 97 nm, 130 nm, 162 nm, 195 nm, 227 nm. Most pores smaller than 97 nm did not quite follow this pattern (although some did, measuring around 33 nm and 65 nm). This pattern suggests that individual pores reach a maximum of ~32.5 nm in diameter, but can fuse together, as observed in images (e.g., Fig. 5 - panel 1), giving rise to larger pores measuring multiples of this diameter.

## 4. Discussion

Pore-forming cytolysins (PFCs) are molecules that interact with biological membranes creating channels that permeate cells. They are predominantly proteins or peptides found in both eukaryotes and prokaryotes, and have been involved in a large spectrum of functions [16, 17, 30–32]. *L. amazonensis* pore-forming cytolysin, which we called leishporin, is a type of such molecules. Previously, using osmotic protection (on erythrocytes) and the whole cell patch clamp technique (on macrophages), we were able to demonstrate that *L. amazonensis* promastigotes extracts damage cells by inducing the formation of nonselective pores on the cell membrane [11, 12]. Although the molecule(s) responsible for *L. amazonensis* pore-forming activity remain(s) unidentified, several features of this cytolysin have been characterized by our group [13–15].

A prominent and powerful technique to evaluate topographic profiles of cell surfaces and artificial bilayers is AFM, which has been used to visualize cell-membrane damage caused by various pore-forming molecules. In the present study, using AFM tapping-mode microscopy, we also could visualize lesions formed on erythrocyte and artificial unilamellar membranes treated with cytolytic *L. amazonensis* mExt. In both cells and artificial membranes, two important traits were found in these lesions: their circular shape and their membrane-spanning thickness (4 to 10 nm) [28, 29], features that distinguish typical pores. Similar images of other pores from various origins, from ions to microrganisms, have been visualized by AFM [23, 33–36], although different images of pores with visible molecules encircling their borders have also been visualized [21, 37–40]. The latter are pores made by protein and they are often visualized as structures in which it is noticeable the assembly of proteins monomers, large enough to be visualized, and/or more well-defined border. Since we still do not know the nature of leishporin molecule(s), the pores visualized here may not resemble protein-made pores. Indeed, not only proteins form pores, and some examples are gadolinium [33], graphene [41], lipids [42, 43], peptides [40], and lipopeptides [34]. In fact, the images of these pores are rather similar to those we show here. This and the fact that we have also identified other non-proteic cytolytic molecules in *L. amazonensis* mExt (unpublished results) make us speculate that leishporin may indeed be a molecule other than protein.

Pores were not visualized on erythrocytes or DPPC liposomes treated with hi mExt, which are not lytic both to erythrocytes and liposomes (Fig. 1), showing that they resulted from the action of active leishporin. Another interesting negative control was the DOPC liposomes, shown here to be resistant to lysis mediated by leishporin. The reason why DOPC, as opposed to DPPC, is resistant to leishporin-mediated lysis is still unknown but is probably neither due to differences in cytolysin binding, since both types of liposomes are able to bind leishporin (Castro-Gomes, Frézard & Horta, unpublished results) nor due to a lysis because of lower fluidity of liposomes, since DPPC liposomes are less fluid than DOPC [35, 36, 44, 45]. Accordingly, pores were also not visualized on DOPC liposomes, again showing the correlation between lysis and visualization of the pores.

Previous results obtained from electrophysiological measurements [12] have shown that diameters of the pores formed by mExt were heterogeneous. At the conditions used, measurements of pores conductance and results from osmotic protection indicated a variation from ~1.6 to > 6.1 nm, but no upper size limit was determined at that condition. Here, the diameters of pores formed in both erythrocytes and lipid films were also quite heterogeneous, but much larger than those previously observed. In erythocytes, pores ranged from 32 nm to 125 nm, while in liposomes, they ranged from 25 nm to 227 nm. The differences in sizes are actually not surprising. In fact, our previous results have shown that the diameter of the pores increases with time and with concentration of leishporin. These data suggested that pores result from 1) aggregation or polymerization of single units and that the numbers of subunits that form each pore determine its diameter or 2) coalescence of two or more individual pores, thus generating the heterogeneity of the pore diameters, as measured by conductance and by osmotic protection [12]. The results presented in the present work strongly support this assumption. Thus, the larger diameters observed here can be attributed to the much higher concentration of mExt and longer incubation time used in the present experiments. It is important to understand that the patch-clamp technique is more apt to detect smaller pores and earlier events than visualization by AFM. In fact, the appearance of larger pores disrupts the osmotic balance of the cell and rapidly leads to the breakdown of membrane resistance and/or the rupture of the gigaseal, precluding the required control of the membrane potential. The fact that pores smaller than 25 nm could not be visualized in erythrocytes or lipid films, after 30 min incubation, suggests a cooperative mechanism in which formed pores favor the incorporation of newer molecules or fully assembled pores, making it more likely the enlargement of the pores than forming new structures. Such mechanism would speed up the osmotic disruption. This type of difference in measurements using different techniques has been already reported. For instance, for EspB and EspD, two hemolytic proteins produced by the type III Secretion System from *E.coli,* osmoprotection assays revealed a minimal pore-size of 3-5 nm whilst AFM showed pores ranging between 55-65 nm [45].

It was quite conspicuous that in liposomes we could visualize pores much larger in diameters than in erythrocytes. One possibility is that, liposomes were more overloaded with mExt than erythrocytes, an approach used in order to maximize the probability to visualize the pores. Another possibility is that in liposomes, pores could more easily fuse with each other giving rise to larger pores. About the variation on the depths of the pores, we can speculate that the shallowest pores (4 nm) might not be completely traversed the membrane and the deepest ones could be due to the fact that the films are not totally straight (in fact, some sort of undulations appears in the images) and the AFM probe could be measuring more than the thickness of the membrane (for instance, in cases where the pore would be in the top of the undulation). A remarkable finding was that, in liposomes, we could observe a pattern in diameters of pores. Many pores had invariably the same diameter, all multiples of ~32.5 nm (Fig. 6), suggesting that individual pores reach a maximum of ~32.5 nm in diameter, but can fuse together, creating larger pores with multiples of this diameter. This maximum size could due to some steric hindrance of molecules that constitute the pore, which hampers further pore growth. Why these quantic increments in pore diameters were not observed with pores in erythrocytes is still unknown. This is corroborated by the frequently observed images showing larger lesions, some apparently shapeless but with circular edges (yellow arrows, Figs. 4 and 5 - panel 2) suggesting that several single circular pores have merged together, also corroborating previous results showing that pores increase with time and with leishporin concentration [12]. Images of pores almost fusing together (Fig. 5 - panel 1) are very suggestive of this assumption. It is worth remembering, however that this situation of large pores was probably made possible due to the unusually high concentration of leishporin used during the lytic assay.

In intracellular pathogens, the main roles proposed for PFCs are: 1) nutrient uptake due to membrane permeabilization, 2) induction of apoptosis through different mechanisms, 3) exit of intracellular microorganisms due to rupture of host phagolysosome and plasma-membrane, 4) traversal of membrane barriers, 5) enhancement of phagocytosis and host cell invasion due to induction Ca^2+^ influx [16, 46–49].

In *Leishmania* life cycle, host cell death/rupture passively caused by amastigote overpopulation has been the customary assumption for release of parasites [4–7]. Lately, due to leishporin activity at both acidic and neutral pH, we have proposed that, it may actively damage macrophage phagolysosomal and plasma membranes, causing amastigotes escape [11–17]. Pore formation has already been shown to participate in the exit of several other microorganisms from host cells, and many PFCs from microorganism origin have been implicated in this process. These are the cases of *Listeria monocytogenes* [50, 51] *Legionella pneumophila* [52], *Toxoplasma gondii* [53] and some *Plasmodium* species [48, 49, 54].

The capacity to cause cell damage on their host cells has been reported as an important feature to provoke the intake of intracellular parasites by host cells. One of the most important consequences of pore-formation is the passage of ions through the plasma membrane, which triggers several cellular events potentially involved with cell invasion and/or evasion by intracellular parasites. An essential ion involved in these processes is calcium. Calcium influx through membrane wounds is known to trigger the process of plasma membrane repair, a mechanism of cell survival that relies on calcium-dependent exocytosis of lysosomes and removal of lesions by endocytosis. This mechanism has been shown to be induced and subverted by the intracellular parasite *T.cruzi,* and pore formation has been shown to facilitate its entry in host cells [55, 56]. *T.cruzi* is phylogenetically correlated with *Leishmania* and produces a cytolytic molecule that shares biochemical characteristics with leishporin [12], which could also perform a similar function. As a matter of fact, unpublished data show that *L. amazonensis* can also invade non-phagocytic cells using apparently the same calcium-dependent exocytosis of lysosomes mechanisms used by *T. cruzi* to invade cells and leishporin is a cytolysin that could facilitate parasite invasion.

The identification and characterization of the molecules responsible for *Leishmania* cytolytic activity is an essential step to provide targets that could limit, reduce or block cell invasion or exit, and we are currently working on this subject. The results presented here provide inequivocal visual evidence of the pore-formation mechanism of leishporin-mediated lysis and, together with published data on the function of the pore-forming cytolysins on the exit/invasion of other intracellular parasites, reinforce the potential role of leishporin in the pathogenesis of leishmaniasis.

## Acknowledgments

We thank Elimar Faria for the valuable technical assistance.

## References

1. Bañuls AL, Hide M & Prugnolle F. *Leishmania* and the Leishmaniases: a parasite genetic update and advances in taxonomy, epidemiology and pathogenicity in humans. Adv Parasitol 2007; 64, 1–109.

2. WHO Expert Committee. Control of the Leishmaniases. Geneva, Switzerland: WHO Press; 2010.

3. Kaye P & Scott P. Leishmaniasis: complexity at the hostpathogen interface. Nat Rev Microbiol 2011; 9, 604–615.

4. Bray RS & Alexander J. *Leishmania* and the macrophage, in: W. Peters, R. Killick-Kendrich (Eds), The Leishmaniasis in Biology and Medicine, Biology and epidemiology, London, Academic Press, 1987; vol. 1, pp 211–233.

5. Wilson ME & Pearson RD. Immunology of leishmaniasis, in: D.J. Wyler (Ed), Modern Parasite Biology, Freeman, New York; 1990. pp. 200–221.

6. Liew FY & O’Donnel CA. Immunology of leishmaniasis, Adv Parasitol 1993; 32, 161–259.

7. Handman E. Cell biology of *Leishmania*. Adv Parasitol 1999; 44, 1–39.

8. Rittig MG & Bogdan C. *Leishmania*-host cell interaction: complexities and alternative views. Parasitol Today 2000; 6, 292–297.

9. Real F, Florentino PT, Reis LC, Ramos-Sanchez EM, Veras PS, Goto H, et al. Cell-to-cell transfer of *Leishmania amazonensis* amastigotes is mediated by immunomodulatory LAMP-rich parasitophorous extrusions. Cell Microbiol 2014; 16, 1549–1564.

10. Noronha FSM, Ramalho-Pinto FJ & Horta MF. Identification of a putative pore-forming hemolysin active at acid pH in *Leishmania amazonensis*. Braz J Med Biol Res 1994; 27, 477–482.

11. Noronha FSM, Ramalho-Pinto FJ & Horta MF. Cytolytic activity in the genus *Leishmania*: involvement of a putative pore-forming protein. Infect Immun 1996; 64, 3975–3982.

12. Noronha FSM, Cruz JS, Beirão PSL & Horta MF. Macrophage damage by *Leishmania amazonensis* cytolysin: evidence of pore formation. Infect Immun 2000; 68, 4578–4584.

13. Castro-Gomes T, Almeida-Campos FR, Calzavara-Silva CE, Silva RA, Frézard FJG & Horta MF. Membrane-binding requirements for the cytolytic activity of *Leishmania amazonensis* leishporin. FEBS Lett 2009; 583, 3209–3214.

14. Almeida-Campos FR, & Horta MF. Proteolytic activation of leishporin: evidence that *Leishmania amazonensis* and *Leishmania guyanensis* have distinct inactive forms. Mol Biochem Parasitol 2000; 111, 363–375.

15. Almeida-Campos FR, Castro-Gomes T, Frézard FJG & Horta MF. Activation of *Leishmania spp* leishporin: evidence that dissociation of an inhibitory molecule not only improves its lipid-binding efficiency but also endows it with the ability to form pores. Parasitol Res 2013; 112, 3305–3314.

16. Almeida-Campos FR, Noronha FSM & Horta MF. The multitalented pore-forming proteins of intracellular pathogens. Microb Infect 2002; 4, 741–750.

17. Horta MF. Pore-forming proteins in pathogenic protozoan parasites. Trends Microbiol 1997; 5, 363–366.

18. Hybiske K & Stephens RS. Exit strategies of intracellular pathogens. Nat Rev Microbiol 2008; 6, 99–110.

19. Roiko MS & Carruthers VB. (2009) New roles for perforins and proteases in apicomplexan egress. Cell Microbiol 2008; 11, 1444–1452.

20. Ruan Y, Rezelj S, Zavec AB, Anderluh G & Scheuring S. Listeriolysin O membrane damage activity involves arc formation and lineaction - implication for *Listeria monocytogenes* escape from phagocytic vacuole. PLoS Pathog 2016; 12, e1005597.

21. Scheuring S, Rigler P, Borgnia M, Stahlberg H, Müller DJ, Agre P et al. High resolution AFM topographs of the *Escherichia coli* water channel acquaporin Z. EMBO J 1999; 18, 4981–4987.

22. Czajkowsky DM, Hotze EM, Shao Z & Tweten RK. Vertical collapse of a cytolysin prepore moves its transmembrane beta-hairpins to the membrane. EMBO J 2004; 23, 3206–3215.

23. Garcia Sáez AJ, Chiantia S, Salgado J & Schwille P. Pore formation by a bax-derived peptide: effect on the line tension of the membrane probed by AFM. Biophys J 2007; 93, 103–112.

24. Mari SA, Köster S, Bippes CA, Yildiz Ö, Kühlbrandt W & Muller DJ. pH-induced conformational change of the β-barrel-forming protein OmpG reconstituted into native *E. coli* lipids. J Mol Biol 2010; 396, 610–616.

25. Chalmeau J, Monina N, Shin J, Vieu C & Noireaux V. α-Hemolysin pore formation into a supported phospholipid bilayer using cell-free expression. Biochim Biophys Acta 2011; 1808, 271–278.

26. Puntheeranurak T, Stroh C, Zhu R, Angsuthanasombat C & Hinterdorfer P. Structure and distribution of the *Bacillus thuringiensis* Cry4Ba toxin in lipid membranes. Ultramicroscopy 2005; 105, 115–24.

27. Richter RP, Bérat R & Brisson AR. Formation of solid-supported lipid bilayers: an integrated view. Langmuir 2006; 22, 3497–3505.

28. Kuchel PW & Ralston GB. Theory and Problems of Biochemistry, Schaum’s Outline/McGraw-Hill; 1988.

29. Hine R. Membrane, in J. Daitith (ed.), The Facts on File Dictionary of Biology, 3rd ed., Checkmark, New York; 1999.

30. Bernheimer AW & Rudy B. Interactions between membranes and cytolytic peptides. Biochim Biophys Acta 1986; 864, 123–141.

31. Ojcius DM & Young JD-E. A role for pore-forming proteins in the pathogenesis by parasites? Parasitol Today 1990; 6, 163–165.

32. Molmeret M & Abu Kwaik Y. How does *Legionella pneumophila* exit the host cell? Trends Microbiol 2002; 10, 258–260.

33. Cheng Y, Liu M, Li R, Wang C, Bai C & Wang K. Gadolinium induces domain and pore formation of human erythrocyte membrane: an atomic force microscopic study. Biochim Biophys Acta 1999; 1421, 249–260.

34. Eeman M, Berquand A, Dufrêne YF, Paquot M, Dufour S & Deleu M. Penetration of surfactin into phospholipid monolayers: nanoscale interfacial organization. Langmuir 2006; 22, 11337–11345.

35. Maherani B, Arab-Tehrany E, Kheirolomoom A, Cleymand F & Linder M. Influence of lipid composition of nanoliposomes encapsulating natural dipeptide antioxidant L-carnosine. Food Chem 2012; 134, 632–640.

36. Khadka NK, Teng P, Cai J & Pan J. Modulation of lipid membrane structural and mechanical properties by a peptidomimetic derived from reduced amide scaffold. Biochim Biophys Acta 2017; 1859, 734–744.

37. Leung C, Dudkina NV, Lukoyanova N, Hodel AW, Farabella I, Pandurangan AP, et al. Stepwise visualization of membrane pore formation by suilysin, a bacterial cholesterol-dependent cytolysin. Elife 2014; 3, e04247. Erratum in Elife 4 2014; e06740.

38. Pfreundschuh M, Hensen U & Müller DJ. Quantitative imaging of the electrostatic field and potential generated by a transmembrane protein pore at subnanometer resolution. Nano Lett 2013; 13, 5585–5593.

39. Czajkowsky DM, Sun J, Shen Y, & Shao Z. Single molecule compression reveals intra-protein forces drive cytotoxin pore formation. Elife 2015; 4, e08421.

40. Giménez D, Sánchez-Muñoz OL & Salgado J. Direct observation of nanometer-scale pores of melittin in supported lipid monolayers. Langmuir 2015; 31, 3146–3158.

41. Duan G, Zhang Y, Luan B, Weber JK, Zhou RW, Yang Z, et al. Graphene-induced pore formation on cell membranes. Sci Rep 2017; 7, 42767.

42. Tanaka Y, Mashino K, Inoue K & Nojima S. Mechanism of human erythrocyte hemolysis induced by short-chain phosphatidylcholines and lysophosphatidylcholine. J Biochem 1983; 94, 833–840.

43. Jimbo T, Sakuma Y, Urakami N, Ziherl P & Imai M. Role of inverse-cone shape lipids in temperature-controlled self-reproduction of binary vesicles. Biophys J 2016; 110, 1551–1562.

44. Maherani B, Arab-Tehrany E, Kheirolomoom A, Geny D & Linder M. Calcein release behavior from liposomal bilayer: influence of physicochemical/ mechanical/ structural properties of lipids. Biochimie 2013; 95, 2018–2033.

45. Jacquot A, Francius G, Razafitianamaharavo A, Dehghani F, Tamayol A, Linder M, et al. Morphological and physical analysis of natural phospholipids-based biomembranes. PLoS One 2014; 9, e107435

46. Ide T, Laarmann S, Greune L, Schillers H, Oberleithner H, & Schmidt MA. Characterization of translocation pores inserted into plasma membranes by type III-secreted Esp proteins of enteropathogenic *Escherichia coli*. Cell Microbiol 2001; 3, 669–679.

47. Ishino T, Chinzei Y & Yuda M. A *Plasmodium* sporozoite protein with a membrane attack complex domain is required for breaching the liver sinusoidal cell layer prior to hepatocyte infection. Cell Microbiol 2005; 7, 199–208.

48. Deligianni E, Morgan RN, Bertuccini L, Wirth CC, Silmon de Monerri NC, Spanos L, et al. A perforin-like protein mediates disruption of the erythrocyte membrane during egress of *Plasmodium berghei* male gametocytes. Cell Microbiol 2013; 15, 1438–1455.

49. Wirth CC, Glushakova S, Scheuermayer M, Repnik U, Garg S, Schaack D, et al. Perforin-like protein PPLP2 permeabilizes the red blood cell membrane during egress of *Plasmodium falciparum* gametocytes. Cell Microbiol 2014; 16, 709–733.50.

50. Bielecki J, Youngman P, Connelly P & Portnoy DA. *Bacillus subtilis expressing a* haemolysin gene from *Listeria monocytogenes* can grow in mammalian cells. Nature 1990; 345, 175–176.

51. Schnupf P & Portnoy DA. Listeriolysin O: a phagosome-specific lysin. Microb Infect 2007; 9, 1176–1187.

52. Alli OA, Gao LY, Pedersen LL, Zink S, Radulic M, Doric M et al. Temporal pore formation-mediated egress from macrophages and alveolar epithelial cells by *Legionella pneumophila*. Infect Immun 2000; 68, 6431–6440.

53. Kafsack BF, Pena JD, Coppens I, Ravindran S, Boothroyd JC & Carruthers VB. Rapid membrane disruption by a perforin-like protein facilitates parasite exit froom host cells. Science 2009; 323, 530–533.

54. Abkarian M, Massiera G, Berry L, Roque M & Braun-Breton C. A novel mechanism for egress of malarial parasites from red blood cells. Blood 2011; 117, 4118–4124.

55. Fernandes MC, Cortez M, Flannery AR, Tam C, Mortara RA, & Andrews NW. *Trypanosoma cruzi* subversts the sphingomyelinase-mediated plasma membrane repair pathway for cell invasion. J Exp Med 2011; 208, 909–921

56. Fernandes MC & Andrews NW. Host cell invasion by *Trypanosoma cruzi: a unique* strategy that promotes persistence. FEMS Microbiol Rev 2012; 36, 734–747.

